# The importance of litter quality for the role of arbuscular mycorrhizal fungi in plant litter decomposition (short communication)

**DOI:** 10.1101/059907

**Authors:** E.F. Leifheit, M.C. Rillig

## Abstract

Arbuscular mycorrhizal fungi (AMF) have been shown to positively and negatively affect plant litter decomposition. The use of litter types with different quality and different observation periods might be responsible for these contradictory results. Therefore, we performed a 10-week laboratory experiment with 7 litter types differing in their C:N ratio, and tested for effects of litter quality and the presence of AMF on litter decomposition. We found that decomposition of plant litter with higher C:N ratios was only beginning and was stimulated by AMF, whereas decomposition of plant litter with lower C:N ratios had already progressed and was decreased by AMF. With this study we show that not only litter quality is important for effects of AMF on litter decomposition, but also the stage of litter decomposition.

---

More than half of the earth’s carbon (C) is stored in terrestrial ecosystems (Le Quéré et al., 2015). Our knowledge of the processes of soil C cycling and potential stabilization of soil carbon is crucial for understanding and predicting global warming (Jobbagy and Jackson, 2000). In this context there is increasing interest in belowground ecosystem processes such as the decomposition of plant litter, which can be strongly influenced by the soil microbial community composition (Averill et al., 2014; Cheng et al., 2012; Cotrufo et al., 2013; Fernandez and Kennedy, 2016).

Arbuscular mycorrhizal fungi (AMF) are widespread organisms that can associate with up to 80 % of the land plant species (Smith and Read, 2008). They have been shown to positively and negatively affect plant litter decomposition: Several studies observed an increase in (easy to decompose) litter decomposition in the presence of AMF, especially under conditions of elevated CO_2_ and nitrogen (N) concentrations (Cheng et al., 2012; Hodge et al., 2001; Koller et al., 2013). The main hypothesis arising from these experiments was that AMF promoted litter decomposition by stimulating the activity of other soil biota, e.g. bacteria and protozoa. However, AMF have also been shown to reduce the decomposition of woody plant litter and of litter in later stages of decomposition, respectively (Leifheit et al., 2015; Verbruggen et al., 2016). Interestingly, studies that observed increased decomposition in the presence of AMF predominantly used easy to decompose materials (‘high quality’ litter or finely ground litter) and studied short-term decomposition, while studies that observed decreasing decomposition in the presence of AMF used either difficult to decompose materials (‘low quality’ litter) or observed longer-term decomposition. Hence, litter quality and the decomposition stage are of crucial importance when studying AMF effects on litter decomposition. In AMF dominated ecosystems both low and high quality litter can be important for nutrient cycling and for the contribution to soil OM, e.g. in AMF-dominated forests and in the case of woody plants and perennials that develop lignified roots in grasslands.

To test the effect of AMF on the decomposition of litter types with different quality we performed a 10- week laboratory experiment with 7 different litter types ranging from herbaceous leaf material to woody fragments. As this was a short-term study, we expected that AMF have a stimulating effect on the decomposition of the added plant litter, especially for the ‘high quality’ substrates.

We used 6 different plant litter types from an experimental field site in Berlin (52° 46′ 71N, 13° 29′ 96E, Germany) and commercially available wood sticks (Meyer & Weigand GmbH, Germany). The OM was analyzed for total C and N concentration and subsequently categorized into three groups of C:N ratios: herbaceous plant leaves were categorized as ‘low’ (C:N ratios 10 – 22), leaves of woody plants were categorized as ‘medium’ (35 – 70) and twigs/pieces of woody plants as ‘high’ (121 – 430) (see Table 1).

**Table 1:** Plant litter types used in the experiment

| Plant litter type | Plant part | C:N ratio | C:N ratio group used in statistical analysis |
| --- | --- | --- | --- |
| *Taraxacum officinale* | Leaves | 10 | Low |
| *Plantago lanceolata* | Leaves | 22 | Low |
| *Ligustrum vulgare* | Leaves | 35 | Medium |
| *Mespilus germanica* | Leaves | 50 | Medium |
| *Acer campestre* | Leaves | 70 | Medium |
| *Symphoricarpos albus* | Twigs | 121 | High |
| *Tilia sp.* | Stem pieces | 430 | High |

150 mg of chopped litter were enclosed in a mesh bag (30 μm pore size) and buried in a conetainer with 375 g of loamy sand, originating from the same field site as the plant litter. For the AMF treatment *Plantago lanceolata* seedlings were inoculated by dipping the roots into a (premixed) powder of a rock flour material (attapulgit clay based powder) containing sterile spores of *Rhizophagus irregularis* (15500 spores g^-1^; Symplanta GmbH & Co. KG) and grown for 10 weeks. Roots of control plants received blank carrier material. Each treatment was replicated 6 times, giving a total number of 84 pots. At harvest plant shoots and mesh bags were removed and dried at 40°C. Plant litter was removed from the mesh bags, cleaned with a brush and the weight loss determined gravimetrically. We analyzed the weight loss of the plant litter using linear regression models and ANOVA with AMF and the C:N ratio group as factors and tested for their interaction in the software R version 3.0.2 (R Core Team, 2013).

Successful root colonization of AM fungal hyphae was confirmed with staining and microscopic magnification using the gridline intersect method (McGonigle et al., 1990; Vierheilig et al., 1998). Hyphal length in the AMF treatments was increased by an average of 1.64 m g^-1^ soil compared to the control (staining and analysis according to Jakobsen et al. (1992)).

The C:N ratio group of the material was of crucial importance for decomposition, and effects were significantly influenced by the presence of AM fungal hyphae (see Fig. 1 and Table 2). We could confirm these findings with an additional ANOVA for the effect of AMF and the C:N ratio as continuous variable (p < 0.0001 for C:N ratio and p < 0.03 for the interaction term of AMF and C:N ratio). Within the C:N ratio groups plant litter with ‘low’ and ‘medium’ C:N ratios generally had higher weight loss, but weight loss was reduced in the presence of AMF compared to OM with ‘high’ C:N ratios, where weight loss was increased in the presence of AMF (suggested by the results of the linear model regression). We did not find the expected effect of AMF stimulating the decomposition of ‘high quality’ litter. However, the overall weight loss in the ‘low’ and ‘medium’ group was between 52 – 77 % and we therefore assume that these litters already were in a later stage of decomposition. In later stages of decomposition competition for nutrients or easily accessible energy sources can be increased and via this mechanism AMF might have been able to decrease the activity of decomposing organisms and thus reduced decomposition of litters with ‘low’ and ‘medium’ C:N ratio (‘high quality’ litters). Accordingly, litters with ‘high’ C:N ratios probably still were in an earlier stage of decomposition at the end of the experiment, given that their weight loss was only 25.8 % on average. Here, C sources and easily available nutrients might not yet have been limiting and AMF could have stimulated the activity of other microorganisms through various mechanisms (see Leifheit et al., 2015) and thus increased the decomposition of ‘low quality’ litter.

**Fig. 1:**
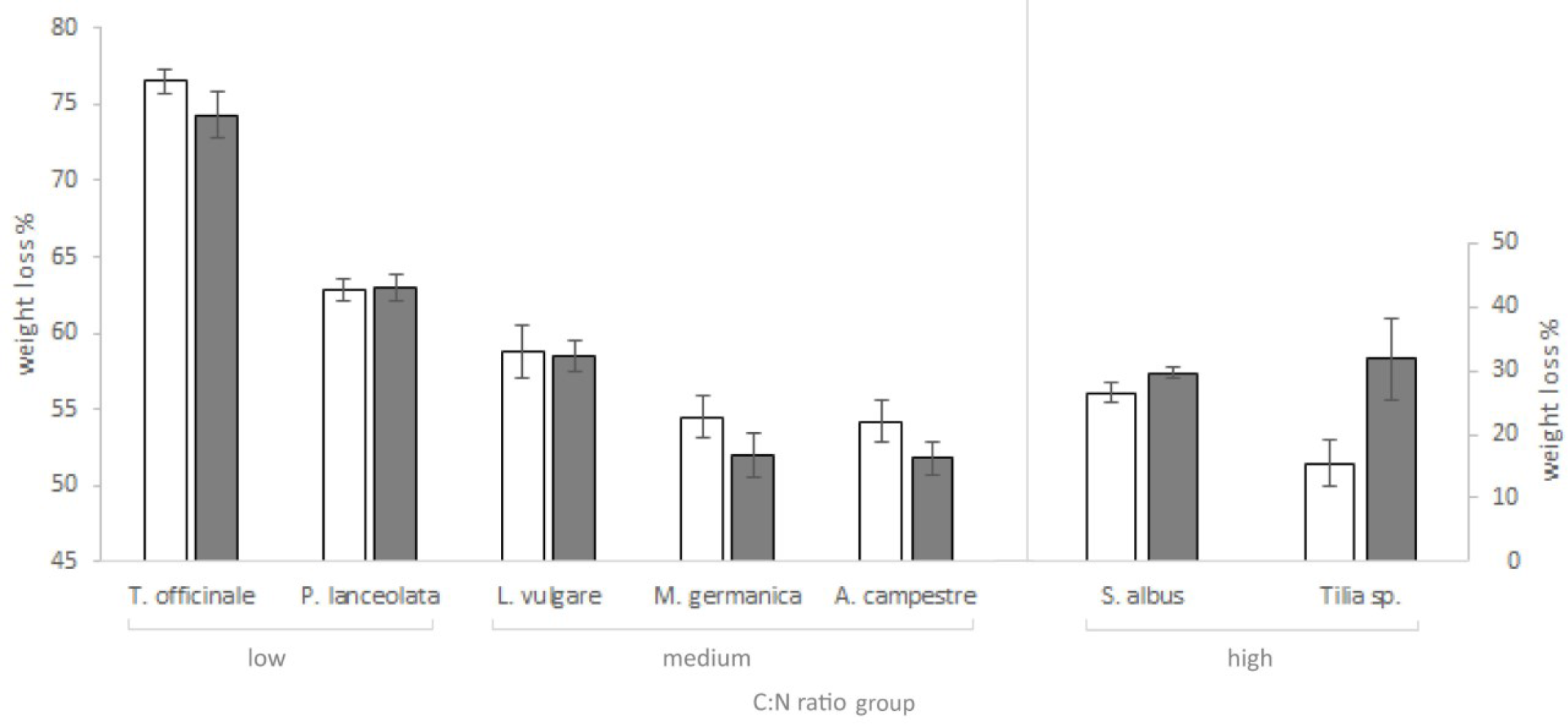
Weight loss (%) of the added plant litter for the different C:N ratio groups (low = 10–22; medium = 35–70; high = 121–430); open bars = without AMF, grey bars = with AMF

**Table 2:** Results of the ANOVA for the presence/absence of AMF and the effect of the C:N ratio groups (‘low’, ‘medium’ and ‘high’, see Experiment design)

|  | p-value |
| --- | --- |
| AMF | 0.77832 |
| C:N ratio group | < 0.0001 |
| AMF:C:N ratio group | < 0.01 |

However, higher litter mass loss or initial faster decomposition in the presence of AMF does not necessarily lead to higher soil C losses. The fate of C in the soil depends - amongst others - on the carbon use efficiency, i.e. the amount of metabolized litter-carbon that is fixed within microbial products in the soil matrix, which could lead to C accumulation in the long-term (Cotrufo et al., 2013; Verbruggen et al., 2013). Furthermore, AMF can contribute to soil C storage through the input of plant photosynthates in combination with stabilization of C within soil aggregates (Rillig and Mummey, 2006; Six et al., 2004; Talbot et al., 2008; Wilson et al., 2009).

With this study we showed that litter quality and the decomposition stage of plant litter is of crucial importance when studying effects of AMF on litter decomposition. Our results confirm previous findings in the literature that AMF can stimulate decomposition in early stages of plant litter decomposition and that they can diminish decomposition in later stages of plant litter decomposition. For future research on AMF effects on litter decomposition it is essential to study the fate of C in the soil, for instance by using labeling techniques.

## Acknowledgements

EFL acknowledges the receipt of a Fellowship from the Dahlem Research School (Germany).

